# Assessment of Metagenomic MinION and Illumina sequencing as an approach for the recovery of whole genome sequences of chikungunya and dengue viruses directly from clinical samples

**DOI:** 10.1101/355560

**Authors:** Liana E. Kafetzopoulou, Kyriakos Efthymiadis, Kuiama Lewandowski, Ant Crook, Dan Carter, Jane Osborne, Emma Aarons, Roger Hewson, Julian A. Hiscox, Miles W. Carroll, Richard Vipond, Steven T. Pullan

## Abstract

The recent global emergence and re-emergence of arboviruses has caused significant human disease. Common vectors, symptoms and geographical distribution make differential diagnosis both important and challenging. We performed metagenomic sequencing using both the Illumina MiSeq and the portable Oxford Nanopore MinION to study the feasibility of whole genome sequencing from clinical samples containing chikungunya or dengue virus, two of the most important arboviruses. Direct metagenomic sequencing of nucleic acid extracts from serum and plasma without viral enrichment allowed for virus and coinfection identification, subtype determination and in the majority of cases elucidated complete or near-complete genomes adequate for phylogenetic analysis. This work demonstrates that metagenomic whole genome sequencing is feasible for over 90% and 80% of chikungunya and dengue virus PCR-positive patient samples respectively. It confirms the feasibility of field metagenomic sequencing for these and likely other RNA viruses, highlighting the applicability of this approach to front-line public health.

## Introduction

Arboviruses are predominantly RNA viruses that replicate in hematophagous (blood-sucking) arthropod vectors such as ticks, mosquitoes and other biting flies to maintain their transmission cycle [1]. These pathogens circulate through sylvatic or enzootic cycles in wild animals causing disease in incidental or dead-end hosts, such as humans, in spill-over events [2]. Human disease outbreaks caused by arboviruses have increased in prevalence over recent decades as ecological factors, human activity and increased vector distribution range have led to the emergence of previously unknown viruses and the re-emergence of viruses previously not thought to be of public health significance [3]. This resurgence has been led by the spread of mosquito-borne arboviruses such as yellow fever (YFV), chikungunya (CHIKV), dengue (DENV), West Nile (WNV), and Zika (ZIKV) viruses across both hemispheres [4]. CHIKV and DENV are of particular global health concern, as they have lost the need for enzootic amplification and have consequently been responsible for extensive epidemics [2].

CHIKV is a single-stranded positive-sense RNA virus of the alphavirus genus which causes the debilitating arthritic disease, chikungunya fever [5]. Originally restricted to regions of Africa and Asia, it has recently spread globally and been designated a serious emerging disease by the WHO [6]. Over the last 15 years, recurring outbreaks of CHIKV appear to have been associated with an increased incidence of more acute presentation of the disease, with an increase in morbidity and possibly mortality [7,8].

DENV is a single-stranded positive-sense RNA virus, member of the flavivirus genus and the most prevalent human arboviral pathogen. DENV fever occurs following infection with one of the four dengue serotypes (DENV1-4). A minority of cases develop severe dengue, with acute hemorrhagic manifestations and multi-organ failure. Despite DENV cases being underreported and misclassified, an increase of 143.1% in dengue fever cases has been estimated between 2005 and 2015 [9]. Approximately 500,000 DENV infected patients require hospitalisation annually [10].

Both CHIKV and DENV are predominantly transmitted to humans via *Aedes* species mosquitoes, particularly *A. aegypti* and *A. albopictus* [11][12], and share clinical presentations of high fever, arthralgia, myalgia, rash and headache. The circulation of CHIKV, DENV (and other arboviruses) in the same geographical locations leads to challenges in differential diagnosis, especially in endemic and epidemic regions in which diagnosis is predominantly symptom-based [13]. Additionally, co-circulation and common vector transmission increase the risk of coinfections. Arboviral co-infection reports have increased in recent years, however, little is known about their clinical presentation and consequences [14–17]. Metagenomic sequencing allows for identification of multiple pathogens within a sample in a non-targeted and unbiased approach. Metagenomic RNA sequencing enables target independent detection and discovery of RNA viruses which has been used for causative agent identification in outbreaks, e.g. Lujo virus in South Africa [18], Bundibugyo ebolavirus in Uganda [19] and novel virus discovery such as the Rhabdovirus associated with hemorrhagic fever in central Africa [20]. In addition to pathogen identification it also provides genomic information for downstream analysis including typing and surveillance. Real-time genomic surveillance was facilitated on-site by the portable Oxford nanopore MinION sequencer during the 2014-2016 EBOV epidemic in West Africa and the ZIKV outbreak in the Americas [21–24] for epidemiological and transmission chain investigations [25]. In both examples, an amplicon sequencing approach was utilised, which has proven extremely useful for known pathogen outbreak samples with low viral titres, but this approach does not allow detection of coinfections or unknown pathogens. Metagenomic approaches on the MinION have in principle sequenced viruses and bacteria from clinical, environmental and vector samples [26–29]. Metagenomic sequencing of CHIKV was also demonstrated in principle on the MinION by Greninger *et al.* in 2015 reporting the detection of CHIKV from a human blood sample [29]. Additionally, metagenomics identified CHIKV coinfections within a ZIKV sample cohort [30], with the high proportion of CHIKV reads identified making it a promising target for the approach.

The resurgence of CHIKV and DENV, and the prevalence of mixed arboviral infections [31–33] create an issue that could ideally be resolved in a single, unbiased and rapid virus detection. In this study we set out to test the feasibility of direct metagenomic sequencing of DENV and CHIKV genomes from a cohort of clinical serum and plasma samples with a representative range of viral loads seen at the time of diagnosis. The objective was to assess the proportion of total nucleic acid present in these samples, that was viral in origin, and to determine the limits of retrieval for whole genome sequencing using both the laboratory based Illumina technology and the portable MinION platform.

## Methods

### Sample Collection and Nucleic Acid Extraction

26 routine diagnostic samples, 9 plasma and 17 serum, were obtained from the Rare and Imported Pathogens Laboratory (RIPL), Public Health England, Porton Down. All had previously tested positive by real-time reverse transcription-PCR (qRT-PCR) for chikungunya or dengue virus. A range of representative clinical Ct values was selected. Total nucleic acid was extracted from 140 μl of each using the QIAamp viral RNA kit (Qiagen) replacing carrier RNA with linear polyacrylamide and eluting in 60 ul, followed by treatment with TURBO DNase (Thermo Fisher Scientific) at 37°C for 30 min. RNA was purified and concentrated to 8 μl using the RNA Clean & Concentrator™-5 kit (Zymo Research)

### Molecular confirmation and quantification

Drosten *et al.* [34] and Edwards *et al. [35]* RT-PCR assays were used for confirmation of DENV and CHIKV respectively. RNA oligomers were used as standards for genome copy quantitation.

### Metagenomic Library Preparation

cDNA was prepared using a Sequence Independent Single Primer Amplification (SISPA) approach adapted from Greninger *et al.* [36]. Reverse transcription and second strand cDNA synthesis were as described. cDNA amplification was performed using AccuTaq LA (Sigma), in which 5μl of cDNA and 1 μl (100 pmol/μl)Primer B (5′-GTTTCCCACTGGAGGATA-3′) were added to a 50 μl reaction, according to manufacturer’s instructions. PCR conditions: 98 °C for 30s; 30 cycles of 94 °C for 15s, 50 °C for 20s, and 68 °C for 5min, followed by 68 °C for 10min. Amplified cDNA was purified using a 1:1 ratio of AMPure XP beads (Beckman Coulter, Brea, CA) and quantified using the Qubit High Sensitivity dsDNA kit (Thermo Fisher).

### MinION Library Preparation and Sequencing

MinION sequencing libraries were prepared using total amplified cDNA of each sample to a maximum of 1 μg. Oxford Nanopore kits SQK-NSK007 or SQK-LSK208 (2D), SQK-LSK308 (1D^2^) and SQK-RBK001 (Rapid) were used and each sample was run individually on the appropriate flow cell (FLO-MIN105, FLO-MIN106 or FLO-MIN107) and run on a 48hr script. Basecalling was performed using Metrichor (ONT) for SQK-NSK007 and SQK-LSK208 or Albacore v1.2 for SQK-LSK308 and SQK-RBK001. Poretools [37] was used to extract fastq files from Metrichor fast5 files.

### Illumina Library Preparation and Sequencing

Nextera XT V2 kit (Illumina) sequencing libraries were prepared using 1.5 ng of amplified cDNA as per manufacturer’s instructions. Samples were multiplexed in batches of a maximum of 16 samples per run and sequenced on a 2 × 150bp paired end Illumina MiSeq run, by Genomics Services Development Unit, Public Health England.

### Data handling

BWA MEM v0.7.15 [38] was used to align reads to the following references (Genbank ID): DENV Serotype 1 (NC_001477.1), DENV Serotype 2 (NC_001474.2), DENV Serotype 3 (NC_001475.2), DENV Serotype 4 (NC_002640.1) and CHIKV (NC_004162.2) using −x ont2d mode for nanopore and MEM defaults for Illumina reads. Samtools v1.4 was used to compute percentage reads mapped and coverage depth. Bedtools v2.26.0 was used to calculate genome coverage at 10x and 20x. Mapping consensus sequences for Illumina were generated using in-house software QuasiBam [39] and for MinION using Racon [40] with pileup function. Taxonomic classification was performed using Kraken (0.10.4-beta)[41] and a locally built database populated with bacterial, viral, and archaeal genomes [42]. De novo assemblies were generated using Spades 3.8.2 [43] in combination with SSPACE Standard v3.0 [44] for Illumina generated sequences and CANU v1.6 [44,45] for Nanopore sequences (settings: corOutCoverage=1000, genomeSize=12000, minReadLength=300, minOverlapLength=50).

## Results & Discussion

### Metagenomic Illumina sequencing of CHIKV and DENV patient samples

To assess the feasibility of direct metagenomic sequencing of CHIKV and DENV genomes from patient serum and plasma we selected a set of positive samples collected during 2016 in RIPL diagnostic laboratories, PHE Porton Down. For each virus, samples were chosen to represent the range of viral titres seen during 2016, based on real-time qRT-PCR Ct value (Figure 1). Ct values were confirmed with a second qRT-PCR and genome copy numbers estimated. CHIKV samples selected ranged from Ct 14.72 to Ct 32.57, corresponding to 10^10^ and 10^5^ genome copies per ml of serum or plasma. DENV samples selected ranged from Ct 16.29 to Ct 31.29, corresponding to 10^9^ and 10^5^ estimated genome copies per ml (Table 1). All samples were subject to DNase digestion prior to reverse-transcription and SISPA. In order to measure the proportion of viral nucleic acid present relative to host/background and assess genome coverage, all samples were sequenced using an Illumina MiSeq, generating between 219,000 and 2,070,000 paired-end reads per sample (Table 1).

**Figure 1.**
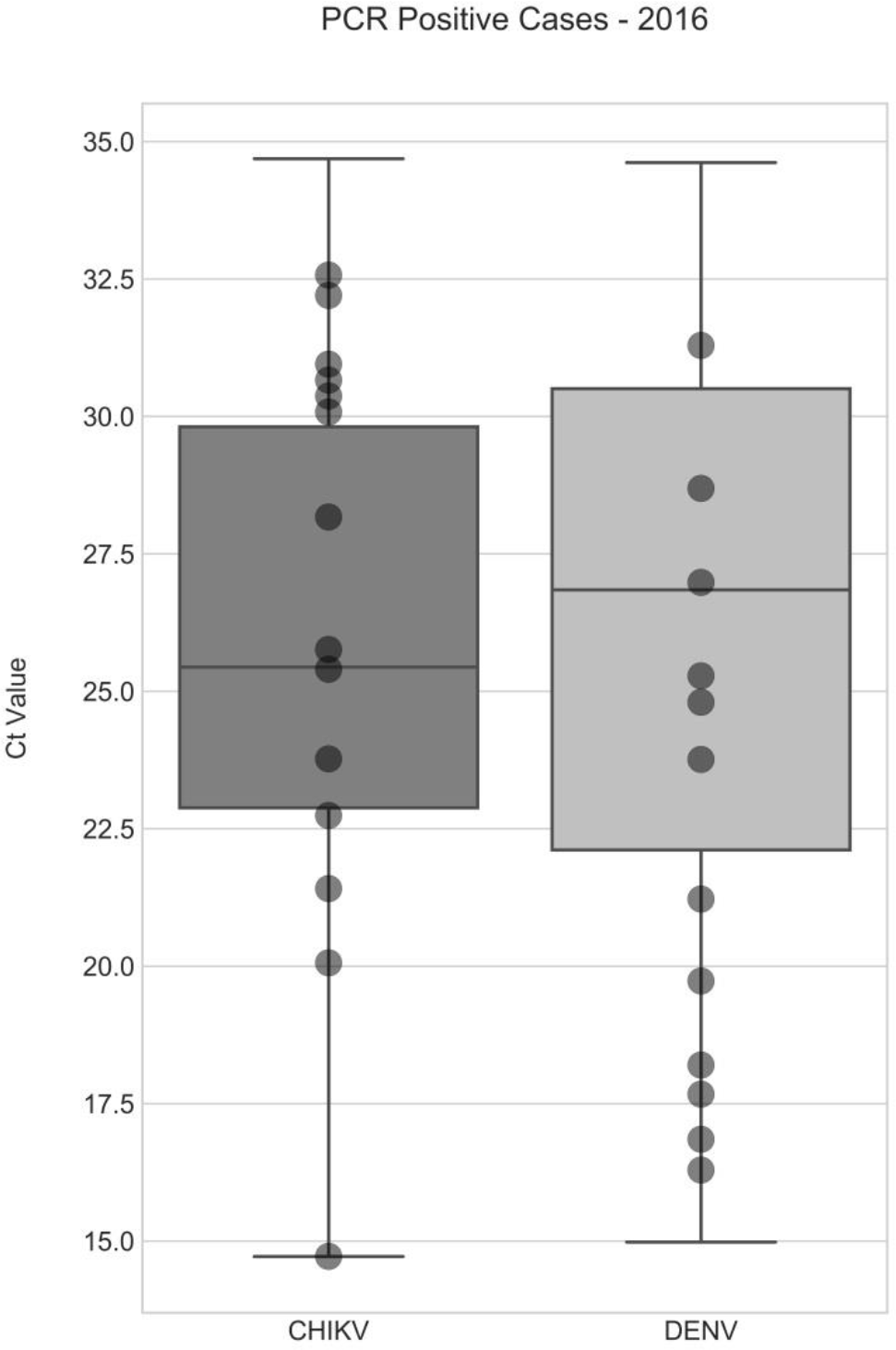
Ct values distribution of RIPL CHIKV and DENV positive samples 2016. A total of 73 samples were positive for CHIKV, and 368 were positive for DENV. Median Ct for CHIKV was 26.1, for DENV it was 26.8. The 14 CHIKV and 12 DENV samples selected for this work are indicated by circles, and were selected to represent a diverse spread of clinical viral loads.

**Table 1.**
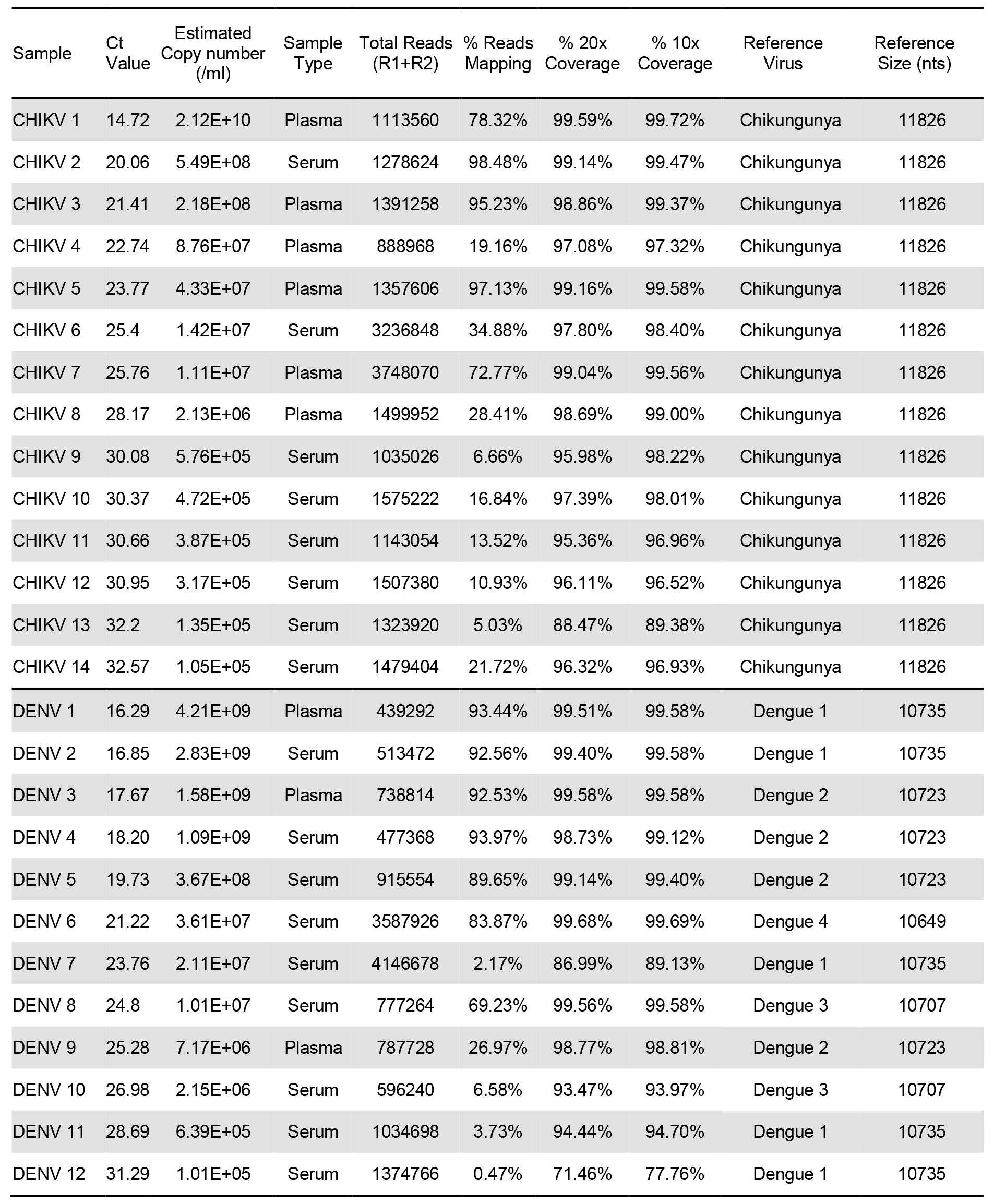
Summary of samples and Illumina mapping data.

### Reference mapping of Illumina data

Reads were mapped to appropriate reference sequences, the percentage of reads mapping and genome coverage at a minimum depth of 20 reads (20x) are shown in Figure 2. The proportion of total reads mapping to the respective viral reference was high for both viruses. In some low Ct samples, over 90% of reads mapped to the viral reference and high proportions were still observed at mid-Ct range. The lowest proportion of reads mapping to the viral reference was 5.03% and 0.47% for CHIKV and DENV respectively (Table 1, Figure 2). The majority of samples returned over 95% genome coverage at 20x and over 98% genome coverage at 10x. Irrespective of lower mapping percentages in high Ct value samples, genome coverage of 88.5% (20x) and 89.4% (10x) for CHIKV and 75.0% (20x) and 77.8% (10x) for DENV was observed. These results clearly show that, due to the significant levels of viral nucleic acid present in CHIKV and DENV clinical samples, relatively modest metagenomic sequencing is capable of elucidating significant portions of viral genome even for samples with Ct values at the higher end of clinical range. A weak correlation was observed between Ct value and proportion of viral reads, with a significant level of variation between samples, likely due to patient-to-patient variability or sample handling during collection, storage and testing. This affected the total level of non-viral host/background nucleic acid present. For example, the two lowest viral titre CHIKV samples (13 & 14) have similar Ct values (32.2 & 32.57) but varied significantly in the proportion of reads that were viral in origin (5.03% & 21.72%). The 5.03% viral reads in CHIKV13 is the lowest for CHIKV, yet still sufficient to generate 88.5% of the CHIKV genome at 20x depth from just ~662,000 paired-end reads. This amount of genomic information is highly informative and further sequencing would very likely increase coverage. Of all diagnostic samples tested in 2016 only 7 of 73 CHIKV samples had a Ct greater than 32.2 (including sample CHIKV14) (Table 1), which suggests that for the majority (>90%) of CHIKV PCR positive samples viral load is sufficient for genome sequencing directly from patient samples without further enrichment beyond a simple DNAse digestion (Figure 1). For the DENV samples, the lowest viral read proportion observed was 0.47% in DENV12, Ct 31.29. This generated 71.5% coverage at 20x depth (increased to 77.8 at 10x depth) from just 687,000 paired end Illumina reads and allowed for DENV serotype identification. Only 62 of 368 DENV cases in 2016 had a higher Ct, predicting that >80% of PCR positive DENV samples have a viral load sufficient for genome sequencing (Figure 1).

**Figure 2.**
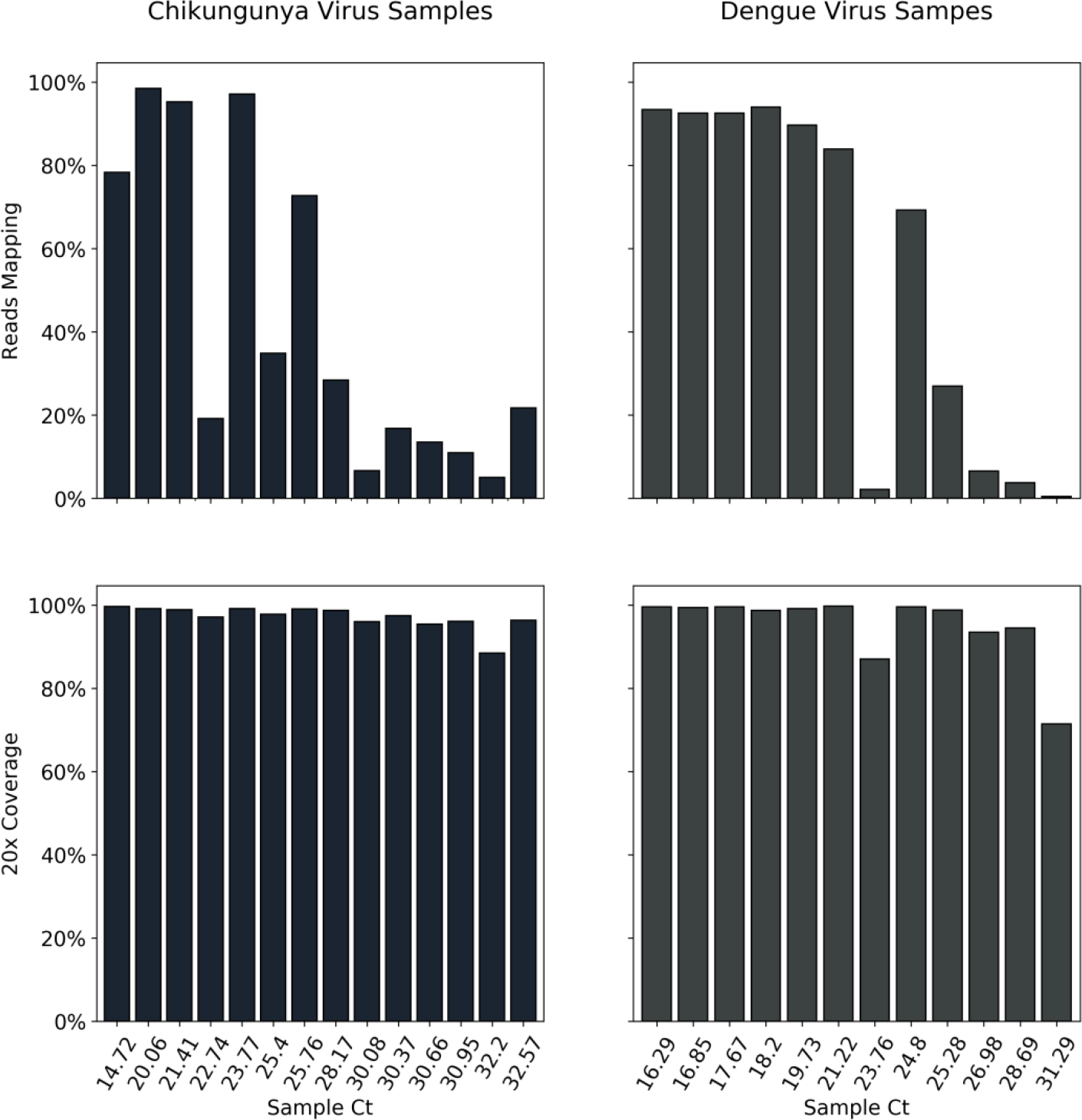
Proportion of viral reads and reference genome coverage per sample. For each sample, identified by Ct value, the percentage of total reads mapping to the appropriate reference sequence, and therefore attributed as viral in origin, is plotted in the upper panel. Lower panels display the percentage of the reference genome sequenced to a minimum depth of 20-fold in the Illumina data generated. See Table 1 for total read numbers.

### Metagenomic MinION sequencing of CHIKV and DENV patient samples

The high yield of viral sequences from clinical CHIKV and DENV samples and recent improvements in data output from the Oxford Nanopore MinION portable sequencing device highlight the exciting prospect of metagenomic MinION viral whole-genome-sequencing, even for lower viral titre samples. To assess this, four samples for each virus from the Illumina sequenced samples were selected to evaluate the performance of the MinION with the metagenomic protocol. Samples were selected with low, mid and high Ct values; CHIKV 1, CHIKV 3, CHIKV 4 and CHIKV 9 (Ct: 14.72, 21.41, 22.74 and 30.08) and DENV 1, DENV 2, DENV 6, DENV 11 (Ct: 16.29, 16.85, 21.22 and 28.69). Total cDNA from the SISPA preparation was used as input for sequencing library preparation and sequenced individually on a single flow cell. Details on total input amounts, sequencing kit and flow cell information for each can be found in Table 2. Total 1D reads generated for each run varied between 203,000 and 3,481,000 per sample and mean read length ranged from 564 to 886 bases (Table 2).

**Table 2.** Summary of samples and Nanopore sequencing.

| Sample | Ct Value | Input (ng) | Sequencing Kit | Flow cell | 1D Total bp | 1D Total Reads | 1D Mean Read Length (nt) | 1D Max Read Length (nt) |
| --- | --- | --- | --- | --- | --- | --- | --- | --- |
| CHIKV 1 | 14.7 | 431.5 | SQK-NSK007 | MIN105 | 1.51E+08 | 267171 | 564 | 92712 |
| CHIKV 3 | 21.4 | 928.8 | SQK-LSK208 | MIN106 | 1.63E+09 | 1891028 | 862 | 99031 |
| CHIKV 4 | 22.7 | 113.4 | SQK-NSK007 | MIN105 | 1.74E+08 | 216493 | 805 | 125387 |
| CHIKV 9 | 30.1 | 212.4 | SQK-LSK208 | MIN106 | 2.12E+09 | 3481358 | 608 | 121711 |
| DENV 1 | 16.3 | 1626.0 | SQK-NSK007 | MIN105 | 2.42E+08 | 284622 | 851 | 115494 |
| DENV 2 | 16.9 | 1626.0 | SQK-NSK007 | MIN105 | 1.55E+08 | 203700 | 760 | 52157 |
| DENV 6 | 21.2 | 475.0 | SQK-LSK208 | MIN106 | 1.22E+09 | 1377721 | 886 | 118733 |
| DENV 11 | 28.7 | 65.8 | SQK-LSK208 | MIN106 | 7.07E+08 | 1111566 | 636 | 119438 |

### Reference mapping of Nanopore data

Extracted 1D reads were mapped to the reference genomes used in the Illumina data analysis. Figure 3 shows percentages of reads mapping to viral reference which were generally concordant with the Illumina data, although a slight decrease is observed across the range of Ct values. In the MinION data the highest mapped read percentages observed were 85.12% and 72.14% for CHIKV 9 and DENV 2 respectively, compared to 95.23% and 92.56% in the Illumina data from the same samples. Whilst in high Ct samples the viral proportion drops to 4.08% for CHIKV 9 and 2.90% for DENV 11, from 6.66% and 3.73% in the Illumina data. Despite the decrease in proportion of mapped viral reads, comparable genome coverage is observed at both 20x and 10x (Figure 3, Table 3) and is even increased compared to Illumina data at lower viral titres, e.g 100% at 20x for CHIKV 9 compared to 95.98% in the Illumina data and 95.25% for the high Ct DENV 11 sample which generated 94.44% coverage from the Illumina data. Differences in precise proportions of viral reads seen are likely due to inter-library variation and do not appear significant. Differences in genome coverage achieved are due to both differences in total reads generated per sample (not normalised between platforms) as well as differences in average individual read length. Average read lengths in Nanopore data ranged from 564 to 886 bp (Table 2).

**Figure 3.**
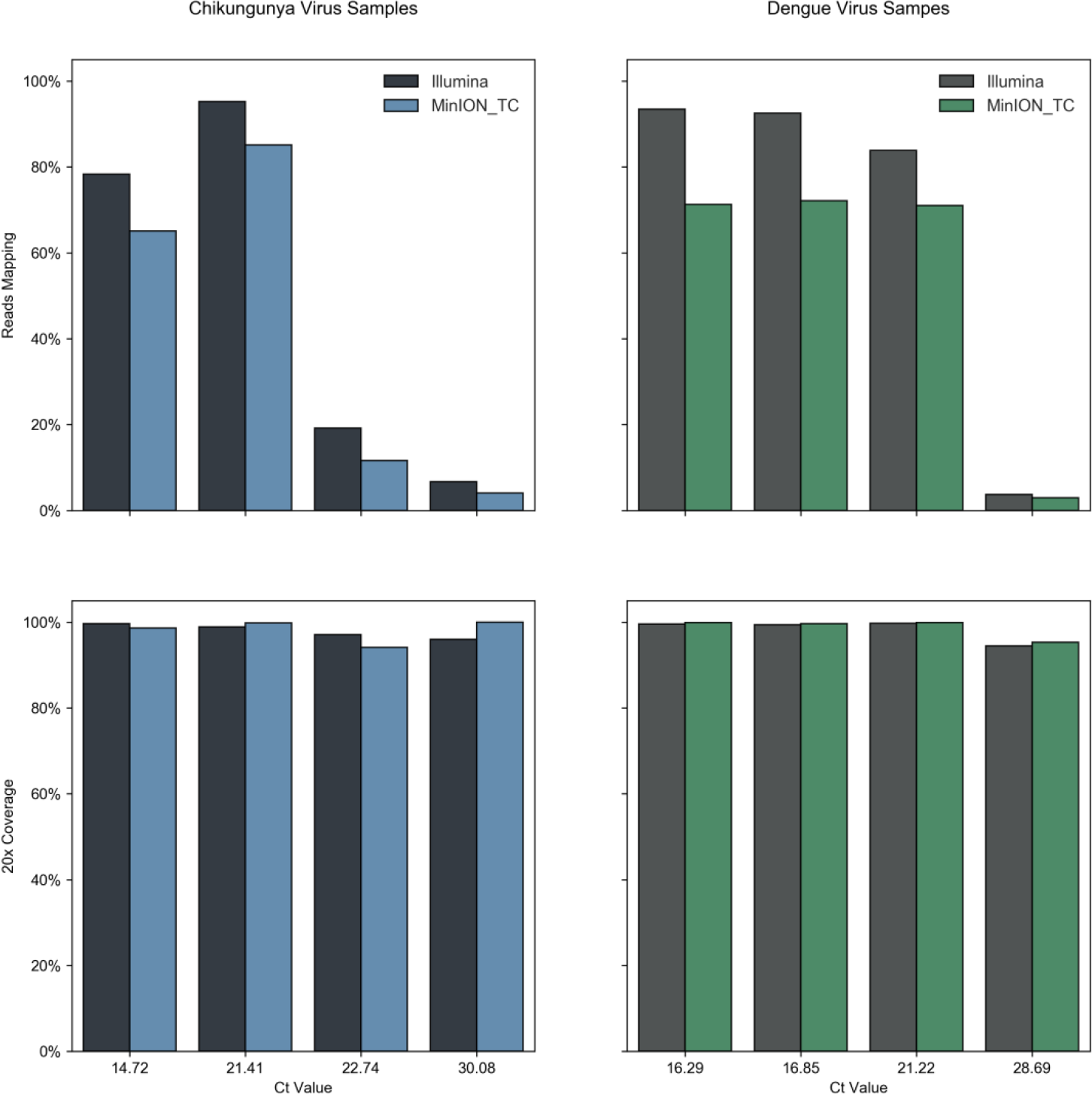
Proportion of viral reads and reference genome coverage comparison. For each sample, identified by Ct value, the percentage of total reads mapping to the appropriate reference sequence, and therefore attributed as viral in origin, is plotted in the upper. Lower panels display the percentage of the reference genome sequenced to a minimum depth of 20-fold in the data generated, in black or grey for the Illumina sequence data, in blue or green for Nanopore data. See Tables 1 & 3 for total read numbers.

**Table 3.** Summary of Nanopore mapping data.

| Sample | Ct Value | Total Reads | % Reads Mapping | % 20x Coverage | 20x Genome Length (nt) | % 10x Coverage | Reference | Reference Size (nt) | Max *de Novo* Contig (nt) |
| --- | --- | --- | --- | --- | --- | --- | --- | --- | --- |
| CHIKV 1 | 14.7 | 267171 | 65.1% | 98.57% | 11658 | 99.2% | Chikungunya | 11826 | 5263 |
| CHIKV 3 | 21.4 | 1891028 | 85.1% | 99.76% | 11798 | 99.9% | Chikungunya | 11826 | 10793 |
| CHIKV 4 | 22.7 | 216493 | 11.6% | 94.11% | 11130 | 97.2% | Chikungunya | 11826 | 4256 |
| CHIKV 9 | 30.08 | 3481358 | 4.08% | 100% | 11826 | 100% | Chikungunya | 11826 | 9860 |
| DENV 1 | 16.3 | 284622 | 71.3% | 99.9% | 10719 | 99.9% | Dengue 1 | 10735 | 8281 |
| DENV 2 | 16.9 | 203700 | 72.1% | 99.6% | 10692 | 99.6% | Dengue 1 | 10735 | 10157 |
| DENV 6 | 21.2 | 1377721 | 71.1% | 99.9% | 10634 | 99.9% | Dengue 4 | 10649 | 7877 |
| DENV 11 | 28.7 | 1111566 | 2.9% | 95.3% | 10226 | 96.3% | Dengue 1 | 10735 | 4699 |

Figure 4 shows coverage depth of reads mapped across the relevant genome for each sample sequenced by both Illumina and Nanopore. Read levels are not normalised thus actual depth is a function of total reads sequenced, but it can clearly be observed that the overall pattern of coverage seen is highly similar for both methods suggesting it is more dependent upon the SISPA methodology than sequencing library preparation. Simple voting consensus sequences generated from each alignment ranged from 99.5% to 99.9% agreement between Illumina and Nanopore data.

**Figure 4.**
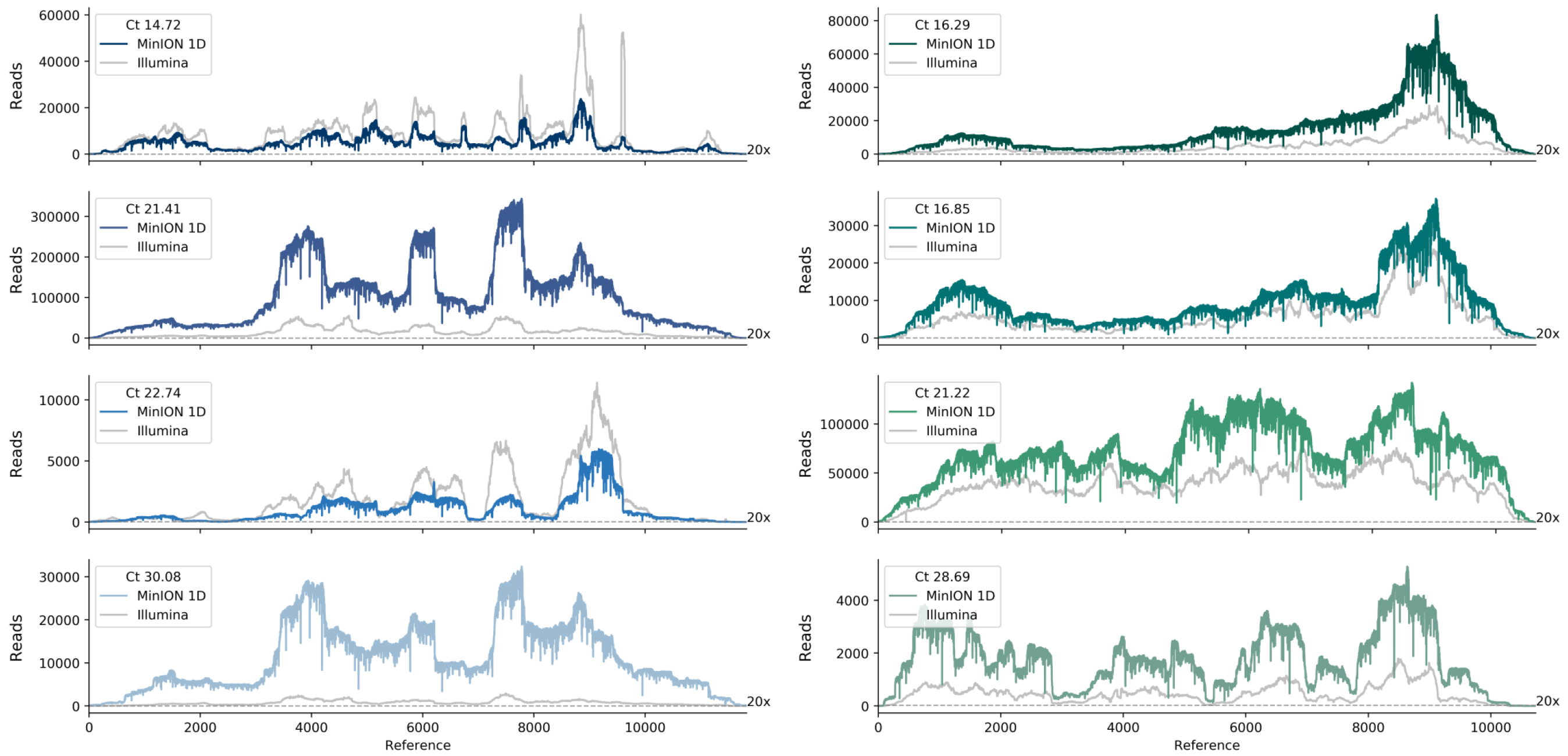
Coverage depth across the CHIKV or DENV genome. Read depth across the genome following reference alignment is shown for all eight samples sequenced by both Nanopore and Illumina method. Illumina coverage is shown in gray, nanopore in blue and green for CHIKV and DENV respectively. Total depth has not been normalised, comparison is to show overall pattern of coverage is highly similar across the methods.

Viral read proportions are generally in agreement with that seen for Illumina sequencing thus it is expected that a similar proportion of qRT-PCR positive patient samples would be suitable for direct metagenomic sequencing on the MinION. Of the samples tested on the MinION, lowest titre samples CHIKV 9 and DENV 11 both generated near complete genome coverage.

### Metagenomic data analysis and co-infection

The data presented clearly demonstrate that direct metagenomic sequencing of serum and plasma samples is a feasible approach for characterisation of CHIKV and DENV with typical viral loads for imported cases sent to PHE’s Rare and Imported Pathogens Laboratory. The true power of a metagenomic approach is unbiased virus identification without prior knowledge of the pathogen. To test the applicability of the metagenomic approach for the data generated, we investigated the target virus detection in each sample by read taxonomic classification using Kraken (Figure 5). The distribution of reads classified as CHIKV, DENV, other viruses, bacteria, archaea/eukaryota and unclassified show a similar pattern for both Illumina and MinION generated reads. The proportion of unclassified reads for each sample increased with Ct value, as the proportion of human origin reads is higher in samples with lower viral load and the human genome is not represented in the Kraken database. A decrease in the percentage of CHIKV and DENV classified reads is observed for MinION data compared to Illumina, due to the lower per read accuracy, but was sufficient to identify the correct predominant virus for all samples.

**Figure 5.**
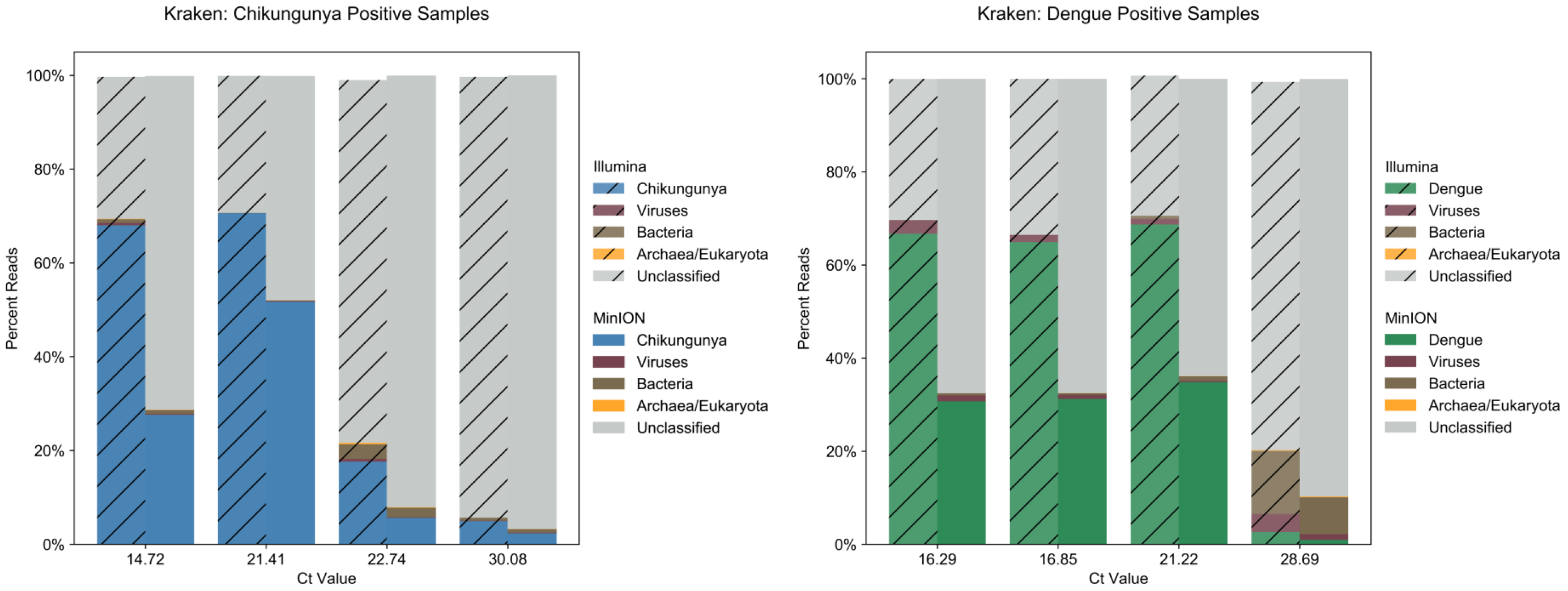
Kraken classification of reads from metagenomic data. Kraken classification distribution comparison for Illumina (cross-hatched) and nanopore data for each sample, for reads grouped as either CHIKV (blue), DENV (green), other viruses (brown), archea/eukaryota (orange), bacteria (brown) or unclassified (grey).

Kraken analysis also allowed for the identification of a DENV co-infection in sample CHIKV 3, the consensus sequence of which (derived via reference mapping) was unique in the sample set, eliminating cross-contamination from the DENV positive samples. For sample CHIKV 3 (CHIKV Ct 21.41) Kraken classified 0.08% of Illumina reads and 0.15% of MinION reads as DENV. Using reference mapping to validate the finding, 0.22% of Illumina reads and 0.43% of MinION reads mapped to DENV-1 reference genome. Genome coverage at 20x of 99.73% and 95.99% was achieved for the primary CHIKV and secondary DENV co-infection respectively, with a single MinION flow cell. This demonstrates the clear benefit of non-targeted approaches for the capacity of libraries generated to detect co-infections even in the presence of another high-titre infection.

### *De novo* assembly

As alternative reference-free approach to read classification *de novo* assembly of the data was attempted using Canu [45] and contig identification by BLASTn. Table 3. lists the longest viral contig length identified in each sample, ranging from 4.2 Kb (36% of reference genome size) to 10.8 Kb (91%) for CHIKV and 4.7 Kb (44%) to 10.1 Kb (95%) for DENV. Identification of the pathogen present without prior knowledge would have therefore been possible for all samples.

### Latest MinION library kits

Initial experiments used analysis of 1D reads from MinION 2D library kit (SQK-LSK208) to successfully identify and genome sequence CHIKV and DENV present in the co-infected CHIKV 3 sample. In order to keep up with the rapidly evolving nanopore chemistries and flowcell developments, we repeated the sequencing utilising the MinION 1D^2^ (SQK-LSK308) and Rapid (SQK-RBK001) kits, currently the most accurate and the fastest library preparation kit available, respectively. The Rapid kit has a library preparation time of just 10 minutes. From this study 74.5% of reads generated with the 1D^2^ kit mapped to CHIKV and 0.37% to DENV, whilst from the Rapid kit the result was 66.26% and 0.29% respectively. A reduction in viral proportion of total reads was observed compared to the 2D kit, which may be due in part to the extended storage time of the original samples prior to retesting. In the case of the 1D^2^ kit, the lower proportion was outweighed by a substantial increase in total data generated per flow cell (5 M vs 1.8 M reads). For the Rapid kit, the lower proportions should be considered in the light of the greatly simplified sample workflow and turnaround-time. Coverage at 20x for CHIKV was above 99% for both kits, and for DENV was 95.04% from the 1D^2^and 81.09% from the Rapid kit (Table 4). Coverage depth across the genome for both viruses can be found in Figure 6, presenting a similar pattern of coverage independent of the kit used. *De novo* assembly of the data (Table 4) produced a CHIKV contig of 10.7, 11.3 and 11.4 Kb for the 2D, 1D^2^ and Rapid libraries respectively and the longest contigs generated for DENV were 7.5, 2.3 and 4.2 Kb. An advantage of the MinION system is the production of data in real-time during the sequencing run. Analysis showed that near-maximum coverage for both viruses was obtained within 30 minutes with the 2D kit, 8 minutes with the 1D^2^ kit and 85 minutes with the Rapid kit (Supplementary Figure 1).

**Table 4.**
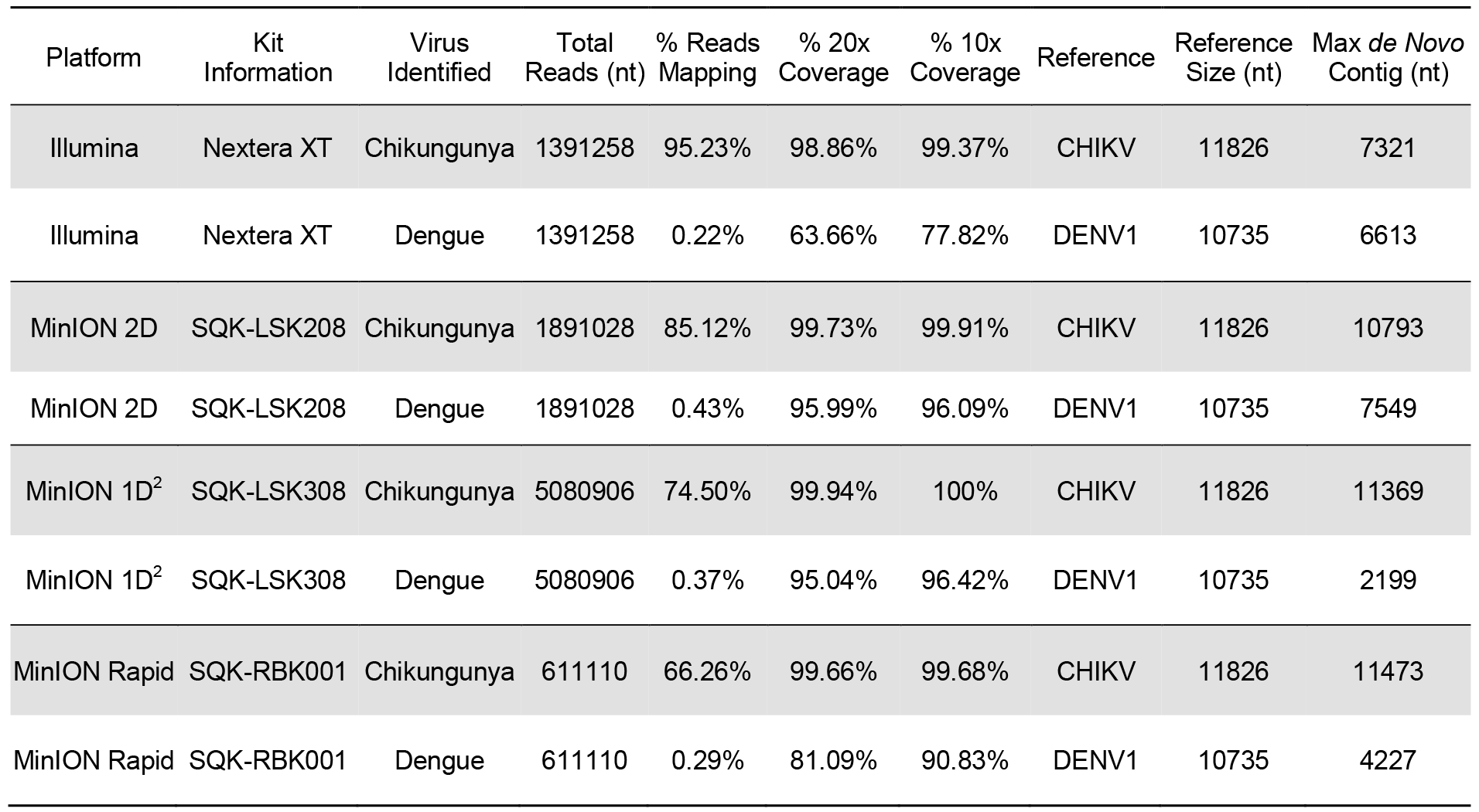
Comparison of Nanopore mapping data across library kits.

**Figure 6.**
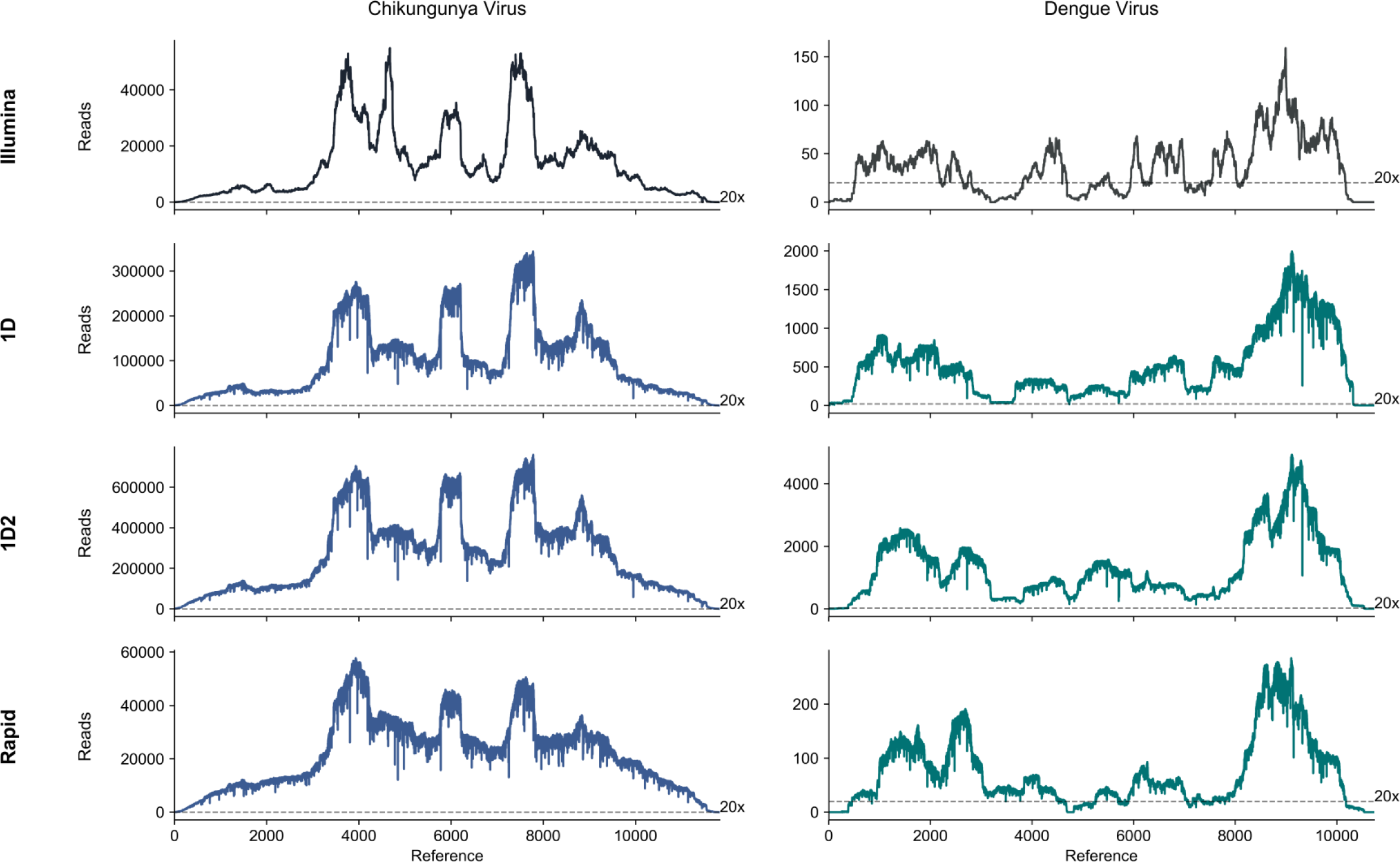
Comparison of genome coverage depth across the CHIKV or DENV genome for the different sequencing library preparation methods. Read depth across both CHIKV and DENV genomes following reference alignment is shown for co-infection sample CHIKV 3, sequenced using for different sequencing library preparation/sequencing methods. Total depth has not been normalised, comparison is to show overall pattern of coverage is highly similar across the methods.

## Conclusions

Globalisation, environmental changes and other human interventions, have increased human risk from emerging and re-emerging viral pathogens [46]. The improvement of diagnostic and genomic sequencing methods, both in speed and detection, will provide an important advantage for disease monitoring and control. We demonstrate that across the clinically relevant range of viral loads an unexpectedly high proportion of reads generated are viral in origin and therefore metagenomic sequencing provides an effective approach for the analysis of CHIKV and DENV genomes directly from the majority of qRT-PCR positive serum and plasma samples, without the need for culture or viral nucleic acid enrichment beyond a simple DNA degradation step. This and recent improvements in the read numbers generated by the MinION, make metagenomic whole genome sequencing feasible in the field. Its use in such a scenario can complement the use of targeted sequencing, as although lower throughput, it has significant additional power in the initial assessment of syndromic outbreaks that lack a clear suspected agent. Once defined, higher throughput targeted methods can be employed alongside. Additionally, the metagenomic approach can detect unexpected causes of disease in the midst of other outbreaks, as well as co-infections. It is ideally suited for the investigation of viral species with high levels of genetic diversity which are difficult to assess using targeted methods, without constant reappraisal. Such issues equally affect diagnostics in non-outbreak scenarios in which the range of RNA viruses to be tested for is both wide and genetically diverse, such as second-line testing of undiagnosed fevers in returning travellers, which therefore may also benefit from a metagenomic approach.

---

**Supplementary Figure 1.**
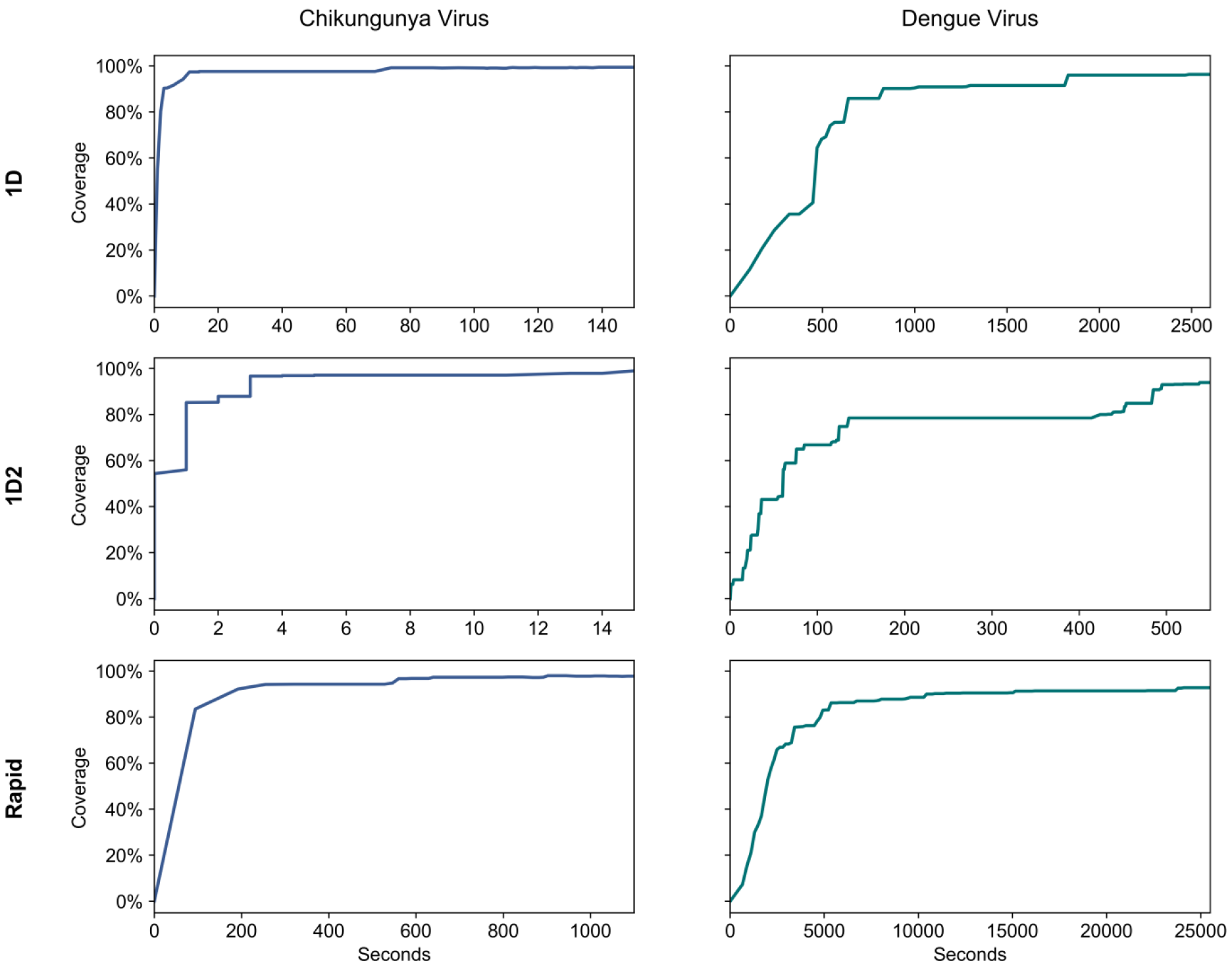
Proportion of genome covered over the course of each sequencing run. The percentage of the CHIKV or DENV genome sequenced plotted over the course of the nanopore sequencing run for each kit version tested.

## Acknowledgements

This work was funded via an NIHR HPRU in Emerging and Zoonotic Infections PhD studentship awarded to L. Kafetzopoulou. The views expressed in this publication are those of the author(s) and not necessarily those of the NHS, the National Institute for Health Research, or the Department of Health. Oxford Nanopore Technologies provided some reagents free of charge and funded author conference attendance.

